# Localization, induction, and cellular effects of tau-phospho-threonine 217

**DOI:** 10.1101/2022.04.18.488657

**Authors:** Binita Rajbanshi, James W. Mandell, George S. Bloom

## Abstract

**Introduction:** Tau phosphorylation at T217 is a promising AD biomarker, but its functional consequences were unknown.

**Methods:** Human brain and cultured mouse neurons were analyzed by immunoblotting and immunofluorescence for total tau, tau_pT217_, tau_pT181_, tau_pT231_ and tau_pS396/pS404_. dSTORM super resolution microscopy was used to localize tau_pT217_ in cultured neurons. EGFP-tau was expressed in fibroblasts as wild type and T217E pseudo-phosphorylated tau, and fluorescence recovery after photobleaching (FRAP) reported tau turnover rates on microtubules.

**Results:** In brain, tau_pT217_ appears in neurons at Braak stages I-II, becomes more prevalent later and co-localizes partially with other phospho-tau epitopes. In cultured neurons tau_pT217_, is increased by extracellular tau oligomers (xcTauOs), and is associated with developing post-synaptic sites. FRAP recovery was fastest for EGFP-tau_T217E_.

**Conclusion:** Tau_pT217_ increases in brain as AD progresses and is induced by xcTauOs. Post-synaptic tau_pT217_ suggests a role for T217 phosphorylation in synapse impairment. T217 phosphorylation reduces tau’s affinity for microtubules.

## 1 INTRODUCTION

The realization that Alzheimer’s disease (AD) comprises a pre-symptomatic phase that might last for more than two decades and that massive, irreversible brain damage has accumulated by the time symptoms are evident has forced a re-evaluation of how to approach the disease clinically. It is now widely accepted that accurate early diagnosis combined with effective interventions that might include drugs and lifestyle adjustments are key to delaying the onset and slowing the progression of AD symptoms, if not preventing symptoms from ever arising. It follows naturally that achieving the tightly linked goals of improved early diagnostics and therapeutics will be aided by a deeper understanding of the molecular mechanisms of AD pathogenesis. While the effort to develop effective therapeutics has not yet yielded any game changing breakthroughs, progress on the diagnostic front has been impressive during the past several years. PET imaging for plaques and tangles is now well established [1–3], and cerebrospinal fluid (CSF) biomarkers have emerged as an alternative approach [4, 5]. PET imaging is sporadically available and costly, though, and obtaining a CSF sample for biomarker analysis requires an invasive procedure in the form of a spinal tap.

Thanks to recent advances in assay sensitivity, it has become possible to substitute blood plasma for CSF to quantify several promising biomarkers for identification of patients at risk of developing AD symptoms years in advance of expected symptom onset. Prime examples of promising plasma biomarkers are phosphorylated forms of the neuron-specific, axon-enriched, microtubule-associated protein, tau, particularly tau_pT181_, tau_pT217_ and tau_pT231_ [6–9]. Though several studies have focused on the localization and functional properties of tau_pT181_ [10] and tau_pT231_ [11], little is known about tau_pT217_ other than its promise as an early fluid-based biomarker for AD, and two reports about its histological distribution in AD brain [12, 13].

As a first step toward understanding functional consequences of tau phosphorylation at T217 we sought to define its distribution *in vivo* by surveying brains of human AD patients at various disease stages and in age-matched, cognitively normal controls, in AD model mouse brains, and in cultured mouse neurons. We also used cultured neurons to determine if extracellular oligomers of amyloid-β (xcAβOs) or tau (xcTauOs) affect the abundance or localization of tau_pT217_. Finally, to determine if tau phosphorylation at T217 affects its affinity for microtubules, we expressed EGFP-tagged wild type (WT) and pseudo-phosphorylated tau in fibroblasts, and used fluorescence recovery after photobleaching (FRAP) to measure turnover rates of microtubule-bound tau.

We found that: 1) tau_pT217_ appears in human brain neurons at Braak stages I-II, and becomes more prevalent at later disease stages in axons, dendrites, neuronal cell bodies, neuropil threads and tangles; 2) in cultured neurons, tau_pT217_ is increased by xcTauOs, but not by xcAβOs, and is associated with developing post-synaptic sites; and 3) pseudo-phosphorylation by a T217E mutation reduces tau’s affinity for microtubules. These results raise the possibility that xcTauOs drive T217 phosphorylation of tau *in vivo*, and that tau_pT217_ contributes to synaptic decline and favors formation of tau-tau interactions at the expense of tau binding to microtubules.

## 2 METHODS

### 2.1 Brain tissue sections

Paraffin-embedded, 5-6 μm thick human cortical brain autopsy sections were obtained from the archives of the Department of Pathology of the University of Virginia, and pertinent clinical information about each donor is shown in Supplementary Table 1. Institutional approval for use of archival autopsy tissue was obtained from the University of Virginia Biorepository and Tissue Research Facility. Classification of disease stages for each donor was performed by ABMS-certified Neuropathologists, James W. Mandell, M.D., Ph.D., and Maria-Beatriz Lopes, M.D., Ph.D., using the “B” (neurofibrillary tangle pathology) part of the ABC staging criteria for AD, in which a score of 0 indicates no detectable tangles, and scores of 1, 2 and 3 respectively correspond to Braak tangle stages I-II, III-IV and V-VI [14].

CVN (APP_SwDI_/NOS2^-/-^) mice [15] were originally obtained from Drs. Michael Vitek and Carol Colton of Duke University, and were maintained as a breeding colony. 5xFAD (APP_SwFlLon_, PSEN1_M146L, L286V_) mouse brain sections were provided by Dr. John Lukens of the University of Virginia Department of Neuroscience. Paraffin-embedded, 5-6 μm thick sagittal sections of brain tissue were cut following trans-cardiac perfusion of 4% paraformaldehyde in PBS of mice that had been deeply anesthetized intraperitoneally with ketamine/xylazine (280/80 mg/kg). Animals were maintained, bred and euthanized in compliance with all policies of the Animal Care and Use Committee of the University of Virginia.

### 2.2 Cultured neurons

As described in our earlier work, primary mouse cortical neuron cultures were prepared from dissected brain cortices of E16-18 WT C57/Bl6 mice and tau knockout (TKO) mice in the same background strain, and were maintained in Neurobasal medium supplemented with B27. For immunofluorescence microscopy, the neurons were grown on 1.5 thickness glass coverslips. All experiments were completed after neurons had been in culture for 10-14 days. Animals were maintained, bred and euthanized in compliance with all policies of the Animal Care and Use Committee of the University of Virginia.

### 2.3 CV-1 cells

CV-1 African green monkey kidney fibroblasts were cultured in Dulbecco’s modified Eagle’s medium supplemented with fetal bovine serum to 10% by volume and 50μg/ml gentamycin.

### 2.4 Immunofluorescence microscopy

Human and mouse brain sections were processed identically. First paraffin was removed with a graded series of xylenes. Next, antigen retrieval was performed by immersing brain slides in a beaker of citrate buffer, and then heating in a microwave at high power for 2.5 minutes followed by low power for 8 minutes. Then, the beaker was slowly cooled to room temperature for 20 minutes, after which it was placed in an ice slurry for 20 minutes. Brain sections were then washed with PBS for 5 minutes and incubated with blocking solution (PBS with 5% bovine serum albumin and 0.1% Triton X-100) for 1 hour, followed by sequential incubations for 2 hours each with primary and secondary antibodies. Multiple washes with PBS followed each antibody step, and immediately prior to being sealed with a coverslip using Fluoromount G (ThermoFisher catalog # 00-4958-02) and Hoechst 33342 (ThermoFisher catalog # 62249), sections were washed for 10 minutes with Autofluorescence Inhibitor Reagent (Millipore catalog # 2160).

An analogous procedure was used for immunofluorescence labeling of cultured neurons following fixation and permeabilization for 5 minutes with methanol at −20° C. In this case, though, primary and secondary antibody incubations were for 30-60 minutes each, and the Autofluorescence Inhibitor Reagent step was omitted.

Dephosphorylation of brain sections and neurons was carried out prior to application of primary antibodies by treating with bovine intestinal mucosa alkaline phosphatase (43 μg/ml, Sigma-Aldrich, Catalog # P7640) overnight at 4° C.

See Supplementary Table 2 for descriptions of all antibodies used for immunofluorescence and western blotting.

An inverted Nikon Eclipse Ti microscope with planapochromatic 10X and 20X dry, and 40X, 60X and 100X oil immersion objectives, a Yokagawa spinning disk confocal head, 4 diode lasers (405, 488, 561 and 640 nm), and a Hamamatsu Flash 4.0 CMOS camera was used for confocal imaging. Quantification of fluorescence micrographs was performed using the [Fiji](https://imagej.net/software/fiji/) software derivative of [ImageJ](https://imagej.nih.gov/ij/), including the Coloc2 plugin for calculating Pearson correlation coefficients.

A dSTORM imaging system (Abbelight; Paris, France) connected to an inverted microscope (DMi8; Leica; Wetzlar, Germany) was used for super resolution microscopy. The microscope was configured with a 63X, NA 1.47 HC planapochromatic oil immersion Leica objective and an EM-CCD camera (Sona 4.2B-6; Andor Technology). Fixed and stained cells were embedded in STORM mounting buffer (Abbelight). Fluorophores were excited using 560 nm (200 mW), and 640 nm (500 mW) lasers. Optimal images were obtained through a combination of corrections and background removal using NEO analysis software (Abbelight).

Pearson correlation coefficients were used for analyzing colocalization and student t tests were used for comparing sample sets. For cultured neurons imaged by dSTORM (Figure 4), a minimum of 50 cells per condition from a total of at least 3 biological repeats were used. Graphs were generated using [GraphPad Prism](https://www.graphpad.com/) 9.0 software. Frequency distribution graphs were generated by first defining each contiguous set of tau_pT217_-positive pixels and PSD95-positive pixels as a single object. Then, for each tau_pT217_-positive object we plotted the shortest distance to a PSD95-positive object, and the shortest distance from each PSD95-positive object to a tau_pT217_-positive object.

**FIGURE 1.**
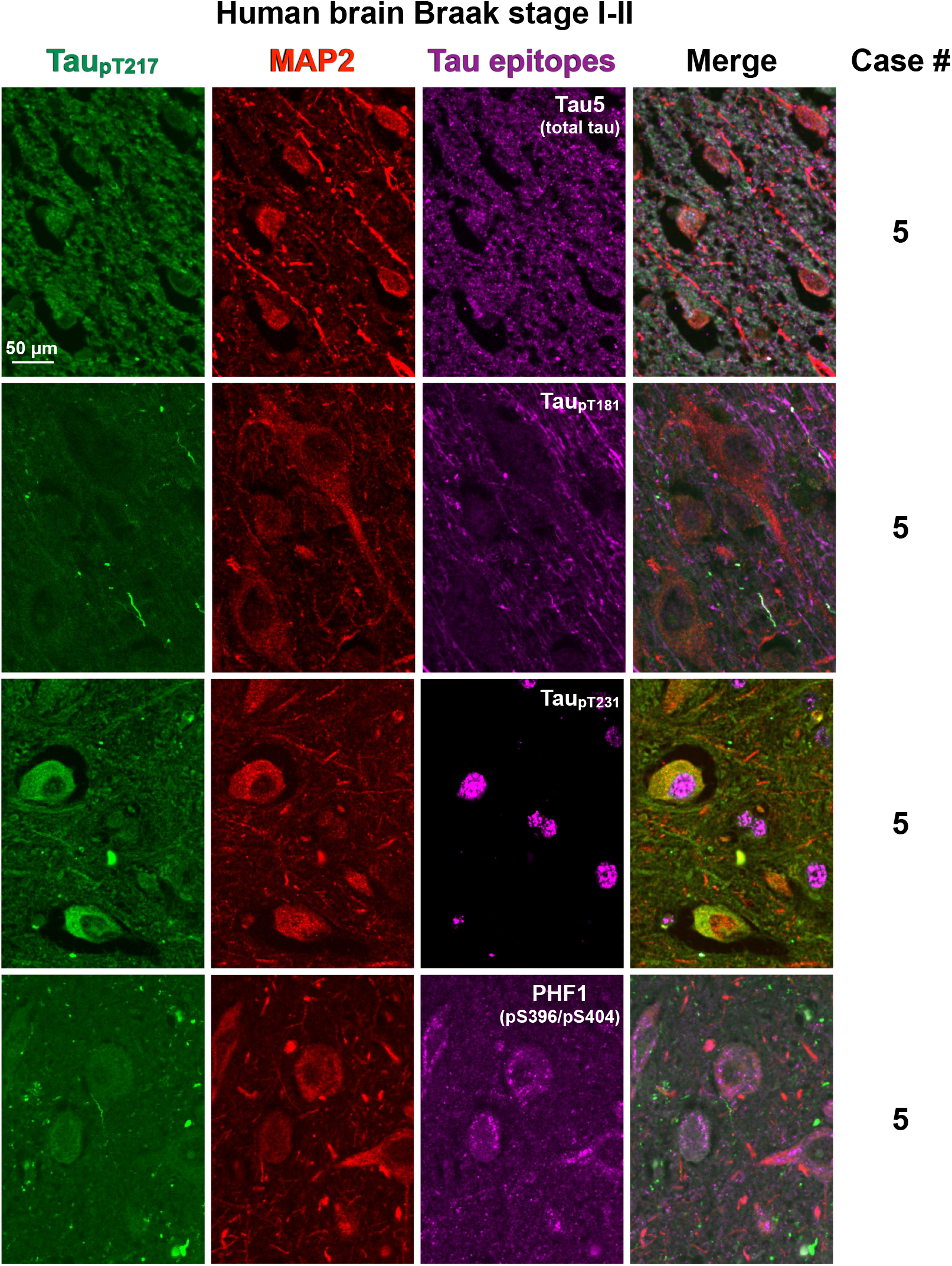
Tau_pT217_ localization in Braak stage I-II human brain cortex. Representative fields of view show ~5 μm thick sections that were triple-labeled with antibodies to tau_pT217_, MAP2, and tau_pT181_, tau_pT231_ or tau_pS396/pS404_, and imaged confocally with a 40X objective. All images illustrate single confocal planes.

**FIGURE 2.**
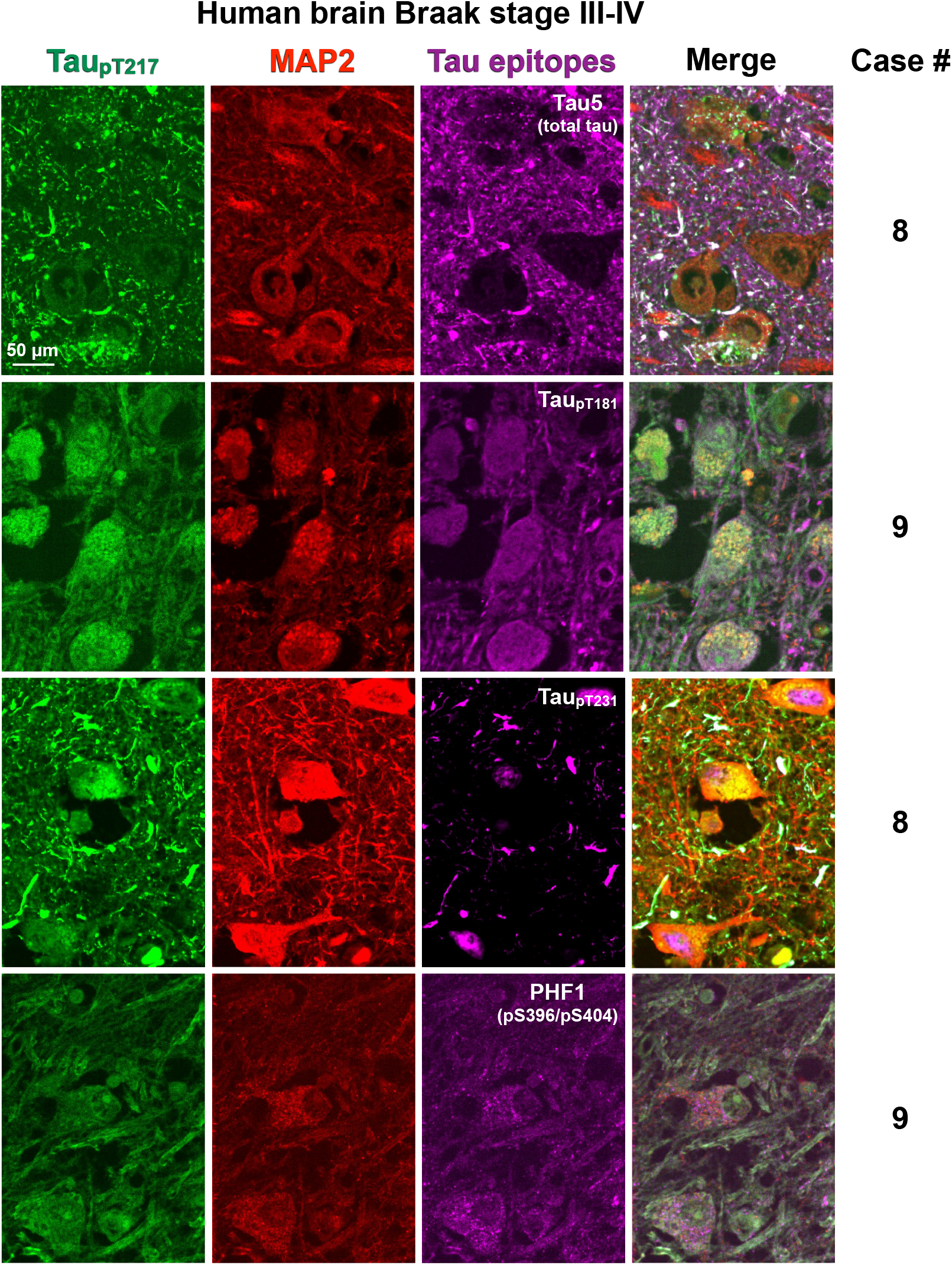
Tau_pT217_ localization in Braak stage III-IV human brain cortex. Representative fields of view show ~5 μm thick sections that were triple-labeled with antibodies to tau_pT217_, MAP2, and tau_pT181_, tau_pT231_ or tau_pS396/pS404_, and imaged confocally with a 40X objective. All images illustrate single confocal planes.

**FIGURE 3.**
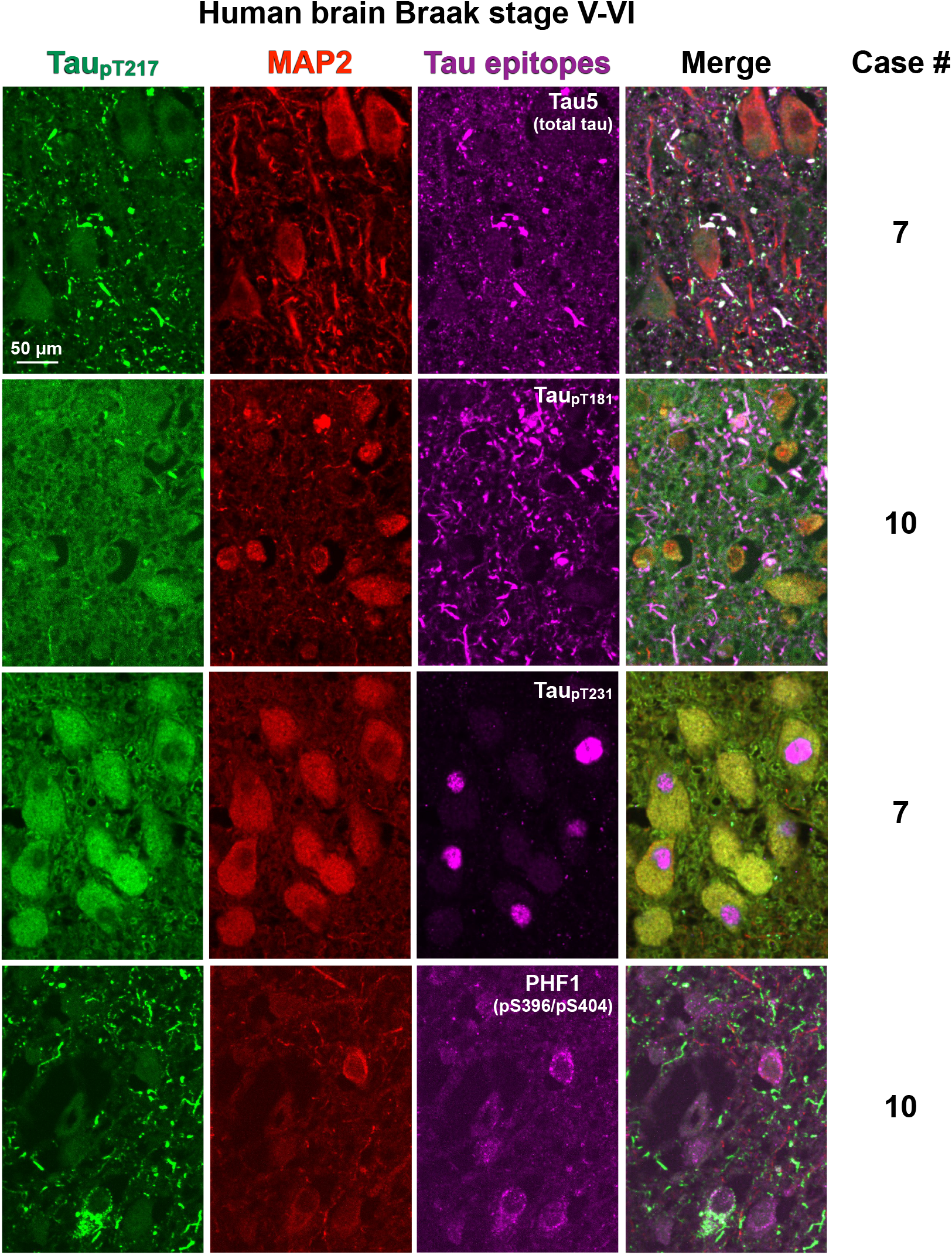
Ttau_pT217_ localization in Braak stage V-VI human brain cortex. Representative fields of view show ~5 μm thick sections that were triple-labeled with antibodies to tau_pT217_, MAP2, and tau_pT181_, tau_pT231_ or tau_pS396/pS404_, and imaged confocally with a 40X objective. All images illustrate single confocal planes.

**FIGURE 4.**
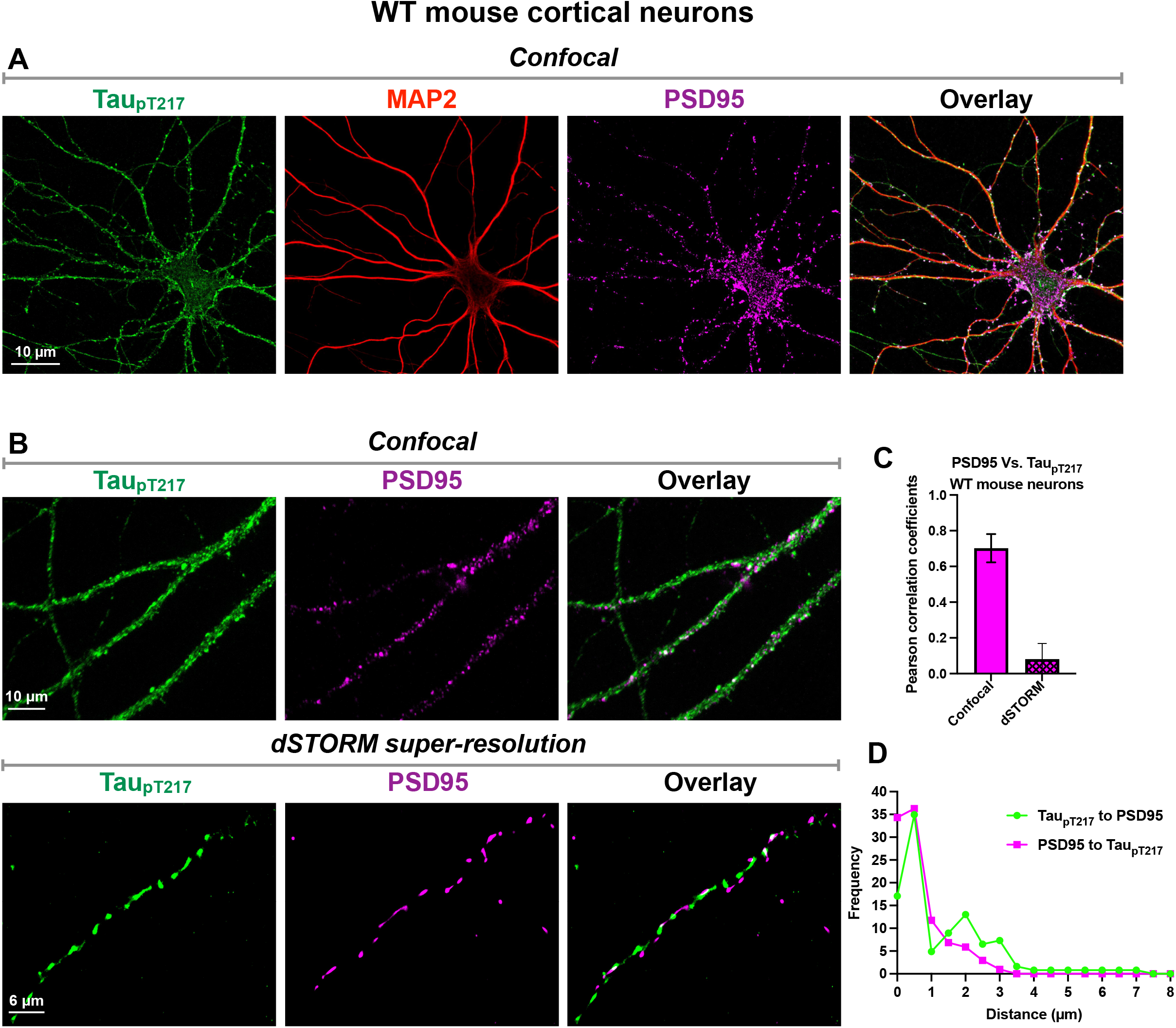
Tau_pT217_ is associated with developing dendritic spines marked by PSD95 in cultured mouse cortical neurons. (A) Single plane confocal images taken using a 60X objective. (B) Single plane confocal (100X objective) and dSTORM super-resolution (63X objective) images. (C) Pearson correlation coefficients for quantifying co-localization of tau_pT217_ and PSD95. (D) Frequency distributions of the nearest tau _pT217_-positive structure to each PSD95-positive structure 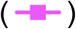 and the nearest PSD95-positive structure to each tau _pT217_-positive structure 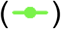. At least 50 cells from each of 3 biological replicates with 12 technical replicates apiece for confocal and 3 technical replicates apiece for dSTORM were used to generate the graphs in panels C and D.

### 2.5 Fluorescence recovery after photobleaching (FRAP) microscopy

pRK5-EGFP-Tau, a mammalian expression vector encoding an EGFP-human 0N4R tau fluorescent fusion protein was obtained from Addgene (catalog # 46904). This plasmid was developed by the lab of Karen Ashe [16], which generously deposited it with Addgene. 0N4R tau was replaced with 2N4R tau after inserting 2N4R tau between BamHI and SalI restriction sites using a cold fusion cloning kit (System Biosciences, Catalog # MC100B-1). A Q5 Site-Directed Mutagenesis Kit (New England BioLabs) was used to convert the WT threonine at position 217 of tau (relative to the largest human CNS tau isoform, 2N4R) to a pseudo-phosphorylated glutamate (T217E). Sequencing of the resulting modified plasmid confirmed the mutation.

WT, T217E (pseudo-phosphorylated) and T217A (non-phosphorylatable) versions of EGFP-tau, and Venus-Tau-Teal [17] were expressed in CV-1 African green monkey kidney fibroblasts using Lipofectamine 3000 (ThermoFisher). The cells were maintained in phenol red-free Dulbecco’s modified Eagle’s medium supplemented with fetal bovine serum to 10% by volume and gentamycin to 50 μg/ml.

FRAP microscopy was performed in three sequential steps (see Figure 6A) using a 488 nm excitation laser at the W.M. Keck Center for Cellular Imaging at the University of Virginia: 1) pre-bleach, in which a region of interest (ROI) was selected and imaged at ~1% power to visualize microtubule-bound tau; 2) photobleach the ROI at 50% power; 3) post-bleach time-lapse imaging for 100 seconds at 1 second intervals at 1% power. Images were captured in each interval on a Zeiss 980 laser scanning microscope with a 63X oil immersion planapochromatic objective. Fluorescence intensity in each ROI was plotted as a function of time before and after photobleaching (see Figures 6B-E).

**FIGURE 5.**
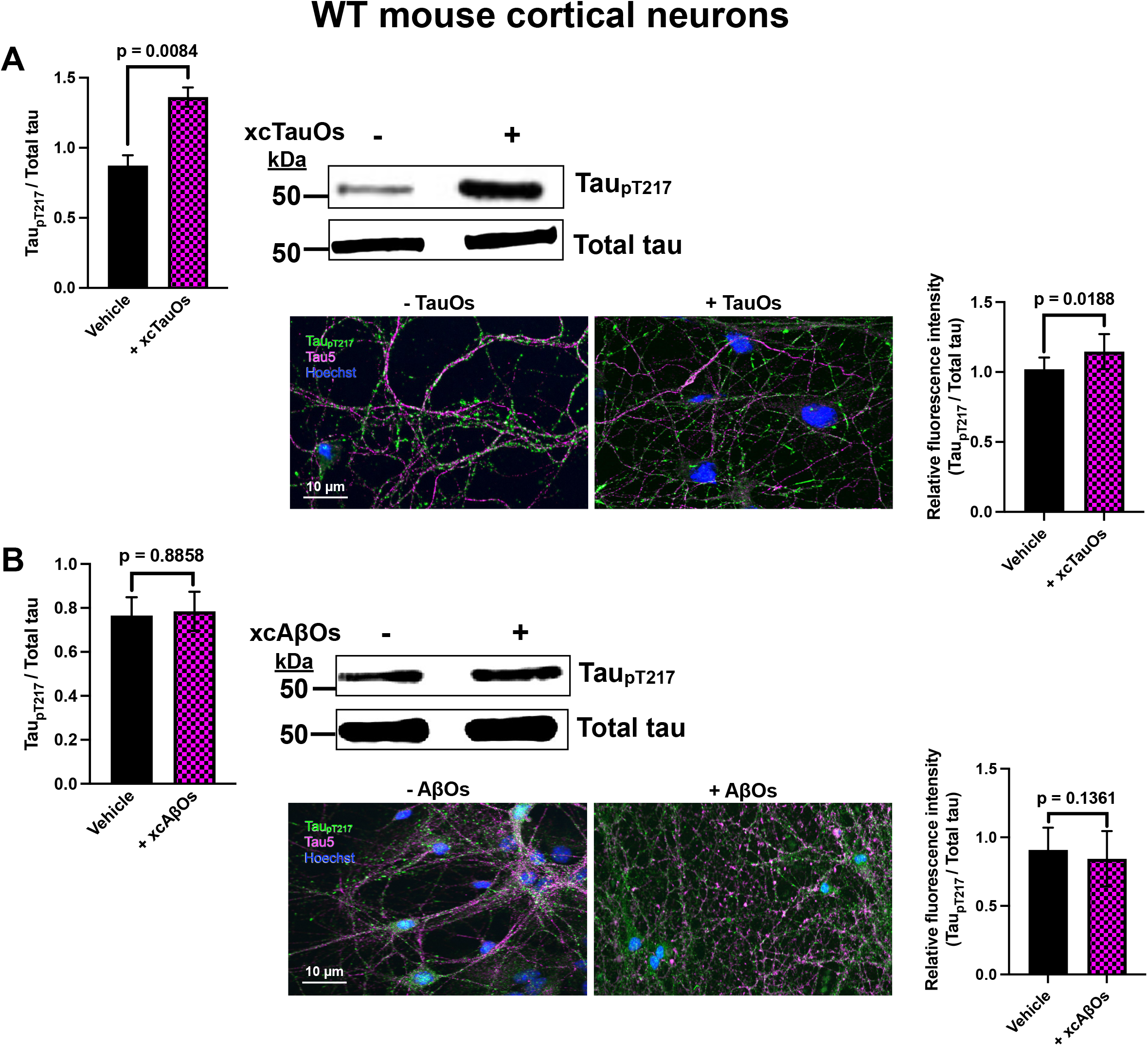
Tau phosphorylation at T217 is stimulated by extracellular tau oligomers (xcTauOs) made from 2N4R human tau (A), but not by extracellular amyloid-β oligomers (xcAβOs) made from Aβ_1-42_ (B). At least 50 cells from each of 3 biological replicates with 4 technical replicates apiece were used to generate the graphs.

**FIGURE 6.**
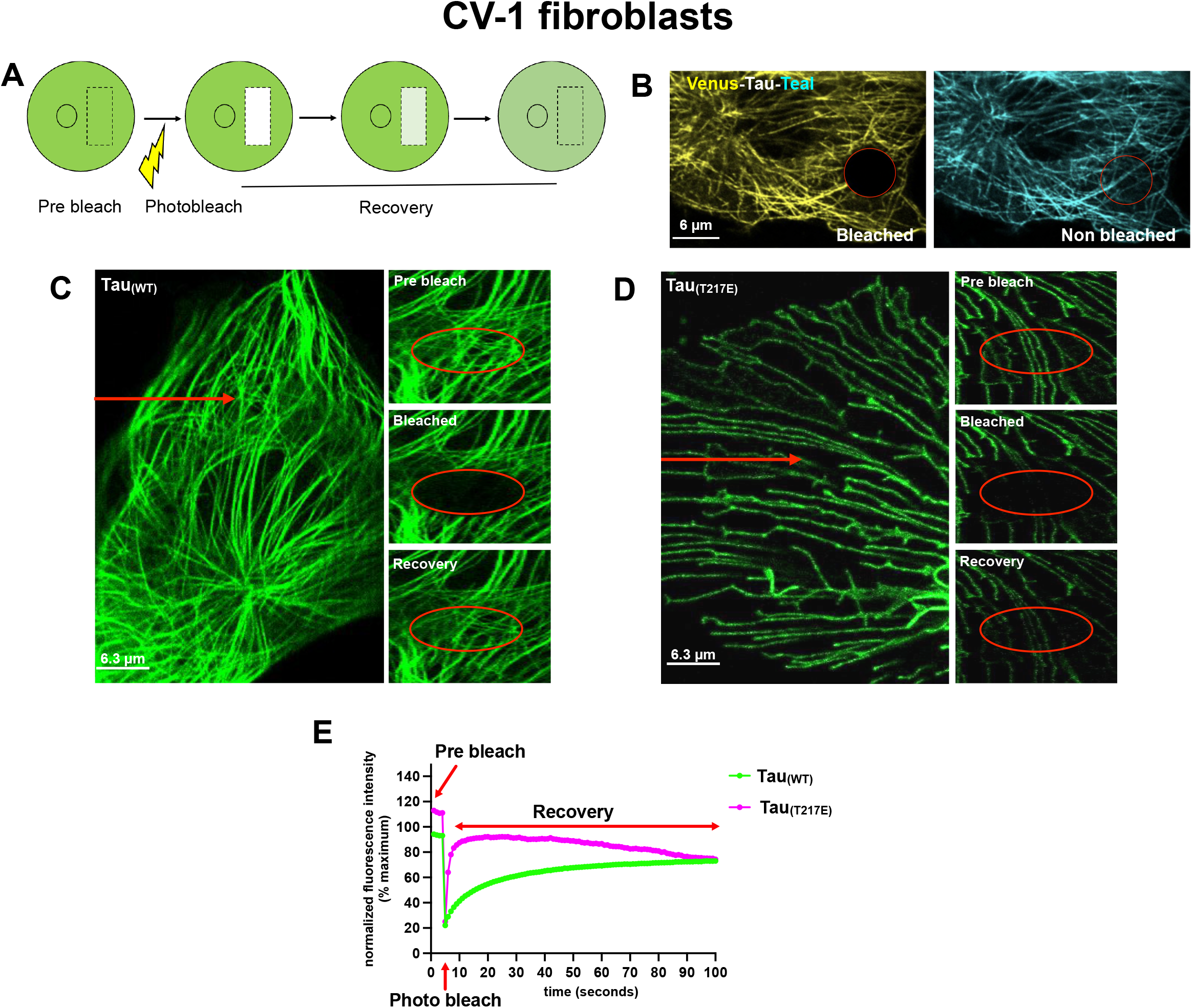
T217E pseudo-phosphorylation reduces tau’s affinity for microtubules. (A) Fluorescence recovery after photobleaching (FRAP) microscopy summary. FRAP images of CV-1 cells transfected to express Venus-Tau_WT_-Teal (B), or WT (C) or T217E (D) human 2N4R tau-EGFP. Note in B that photobleaching of Venus did not remove tau from the microtubules, because microtubules were still visible in the Teal channel. (E) Quantification of FRAP. At least 23 cells from at least 3 biological replicates with 3 technical replicates apiece were used to generate the graphs for both Tau_WT_-EGFP and Tau_T217E_-EGFP.

To normalize the FRAP data (equation 1), the bleached ROI intensity (*F*(*t*)_*ROI*_) was divided by the whole cell intensity for each time point (*F*(*t*)*_cell_*) after subtracting background fluorescence (*F_bkgd_*) at each time point to correct for the loss of fluorescence during the bleach step. The data were then normalized to the pre bleach intensity (*F_i-cell_, F_i-ROI_*) and multiplied by 100 to yield a percentage of initial fluorescence. The resulting normalized data were then averaged for different cells and generated the graphs.

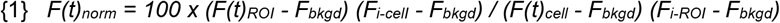

To calculate the mobile fraction of molecules (*Mf*; equation 2) that recover during the time course of the experiment, we used the data obtained from the normalized recovery curves (equation 1) where F_E_, F_0_, and F_i_ are the normalized fluorescence intensities at the end of the experiment, immediately following the bleach, and prior to the bleach, respectively.

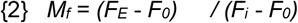

We then calculated immobile fraction of molecules (*IM_f_*) by using equation 3.

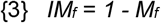

Halftime of recovery (t_1/2_), corresponding to 50% of recovered fluorescence intensity (I_1/2_), was then monitored. To compare the halftimes of a molecule under different experimental conditions, ROI size was held constant.

### 2.6 Extracellular Aβ_1-42_ oligomers (xcAβOs)

Lyophilized, synthetic Aβ_1-42_ (AnaSpec, Catalog # AS20276) was dissolved in 1,1,1,3,3,3-hexafluoro 2-propanol (Sigma-Aldrich) to ~1 mM and evaporated for 4 hours at room temperature. The dried powder was resuspended for 5 minutes at room temperature in 40 μl of 20mM sodium hydroxide and sonicated for 10 minutes in a water bath. To prepare xcAβOs, the monomeric peptide was diluted to 400 μl (100 μM final concentration) in Neurobasal medium (GIBCO) and incubated for 48 hours at 4° C with rocking. This was followed by centrifugation at 14,000 g for 15 minutes to remove fibrils and any other large aggregates that may have been present. Cultured neurons were exposed to 2 μM xcAβOs for 16-18 hours before being processed for immuno-fluorescence or western blotting.

### 2.7 Extracellular tau oligomers (xcTauOs)

A bacterial expression plasmid for human 2N4R tau his-tagged at its C-terminus was generously provided by Dr. Nicholas Kanaan of Michigan State University. The protein was expressed in competent BL21 *E. coli* cells induced with 100 mM IPTG and purified from bacterial extracts using TALON beads (Takara Bio, Catalog # 635502) according to the vendor’s instructions.

Purified tau was diluted to 8 μM in a solution of 100 mM NaCl, 5 mM dithiothreitol, 10 mM HEPES (pH 7.6), 0.1 mM EDTA and 300 μM arachidonic acid (AA; Cayman Chemicals, Catalog # 90010). The protein was then allowed to aggregate for 18 hours at room temperature in the dark. Resulting xcTauOs were diluted into neuron cultures to yield a final total tau concentration of 250 nM. 24 hours later, cultures were processed for immunofluorescence or western blotting.

### 2.8 SDSPAGE and Western blots

Cultured neurons, mouse brains and CV-1 fibroblasts were lysed using N-PER Neuronal Protein Extraction Reagent (ThermoFisher catalog # 87792). Samples were resolved by SDSPAGE using 10% acrylamide/bis-acrylamide gels and transferred to 0.2 μm pore size nitrocellulose (Bio-Rad). Membranes were blocked with Odyssey blocking buffer (LI-COR) and were incubated with primary antibodies and LI-COR secondary IRDye-labeled antibodies diluted into Odyssey blocking buffer. After each antibody step, the blots were washed five times in PBS containing 0.1% Tween 20. The blots were imaged using a BioRad ChemiDoc MP. See Supplementary Table 2 for descriptions of all antibodies used for immunofluorescence and western blotting.

Dephosphorylation of blots was carried out prior to application of primary antibodies by treating with bovine intestinal mucosa alkaline phosphatase (43 – 60 μg/ml, Sigma-Aldrich, Catalog # P7640) overnight.

## 3 RESULTS

### 3.1 The challenge of antibody specificity: presence of the tau_pT217_ epitope in multiple proteins

Throughout the course of this study, the only commercially available antibodies against tau_pT217_ were affinity purified rabbit polyclonal IgG made against a peptide surrounding pT217 in human 2N4R tau. Products fitting this description are sold by multiple vendors, including but not necessarily limited to ThermoFisher, GenScript, Medimabs, and Abcam, the last of which was the source of the anti-tau_pT217_ that we used. Some vendors, including Abcam, list the immunogen as the LPpTPP peptide that is found in the proline-rich region of all human and mouse tau isoforms, and corresponds to amino acids 215-219 in human 2N4R tau, with the threonine at position 217 being phosphorylated. The other vendors do not specify the immunogen in such detail, but it is possible that they all sell the same product.

Because the peptide used to make Abcam anti-tau_pT217_ was so short, we were concerned that it might be present in other proteins, in which case antibody specificity for tau_pT217_ could be problematic. To determine if the immunogen peptide exists in other mouse or human proteins, we used [PhosphoSitePlus](https://www.phosphosite.org/homeAction.action) to search for proteins with documented threonine phosphorylation within an identical LPpTPP peptide. Numerous perfect hits were detected by the search (see Supplementary Table 3), emphasizing the need to seek conditions in which the anti-tau_pT217_ recognizes only tau phosphorylated at T217.

In light of the potential for anti-tau_pT217_ to recognize multiple proteins we established the following criteria that must be met to justify using the antibody for immunolocalization of tau_pT217_ in cultured neurons and brain tissue. On multiplexed western blots of cultured neuron and brain extracts the antibody must: **1)** label one band or multiple closely spaced bands that co-migrate with bands labeled by the Tau5 mouse monoclonal anti-tau, which recognizes an epitope that is found in all tau isoforms and is not subject to any post-translational modifications [18]; **2)** the immunoreactive bands must be sensitive to treatment of blots with alkaline phosphatase prior to incubation with anti-tau_pT217_; and **3)** the anti-tau_pT217_ must not label any bands in extracts of brain tissue or cultured neurons derived from tau knockout (TKO) mice. All of those criteria were met by diluting the antigen affinity purified anti-tau_pT217_ 5000-fold to 200 ng/ml (1.43 nM; Supplementary Figure 1). At concentrations of 1 μg/ml (7.14 nM) or higher, though, the Abcam anti-tau_pT217_ also recognized a prominent ~100 kDa protein that is found in WT and TKO mice, and in human brain (not shown) so that band cannot represent any form of tau. Although we have not determined the identity of the ~100 kDa band, based on its size it might represent Map3k14, which is ubiquitously expressed, and like tau_pT217_ contains the LPpTPP sequence. It is important to note that other anti-tau_pT217_ antibodies might be similarly affected by this specificity issue. Examples of such antibodies include the aforementioned rabbit polyclonals sold by vendors that do not divulge the immunogen sequence, and Eli Lilly and Company’s IBA493, which is being used for AD biomarker research [12] and is not commercially available.

Another condition we established to inspire confidence that anti-tau_pT217_ specifically and exclusively recognizes tau phosphorylated at T217 for localization is to show by immunofluorescence that staining of brain tissue and primary neuron cultures can be eliminated when the cells or tissue are incubated with alkaline phosphatase prior to the primary antibody step. Supplementary Figure 2 illustrates that when anti-tau_pT217_ is used at 200 ng/ml for immunofluorescence, it preferentially labels neurons in primary mouse cortical neuron cultures that also contain glial cells, as well as in CVN mouse and human AD brain tissue. Moreover, pretreatment of the cultured cells or brain tissue sections with alkaline phosphatase abolished staining by anti-tau_pT217_. We thus conclude that using anti-tau_pT217_ at a concentration of 200 ng/ml allows unambiguous detection of tau_pT217_ in brain tissue and cultured neurons by both western blotting and immunofluorescence.

Similarly to the anti-tau_pT217_ antibody, the specificities of mouse anti-tau_pT181_ and mouse anti-tau_pT231_ were tested in primary WT and TKO mouse cortical neurons. At a concentration of 50 ng/ml both anti-tau_pT181_ and anti-tau_pT231_ labeled cultured WT neurons under basal conditions by immunofluorescence (Supplementary Figure 3). When the antibodies were used for western blotting at that concentration, each labeled a single band that co-migrated with a total tau band labeled by Tau5 in WT neuron extracts, but no immunoreactive bands were observed for extracts of TKO neurons (Supplementary Figure 4). We therefore conclude that the anti-tau_pT181_ and anti-tau_pT231_ antibodies were specific for tau phosphorylated at T181 and T231, respectively, under the conditions in which we used them.

### 3.2 Tau_pT217_ localization in human AD and transgenic AD mouse model brain

To study the *in vivo* localization of signature AD phospho-tau epitopes, we performed triple immunofluorescence microscopy of human and murine brain sections using antibodies specific for tau_pT217_ and the dendrite-enriched protein, MAP2, plus tau_pT181_ (AT180 epitope, but detected by a different antibody), tau_pT231_, or tau_pS396/pS404_ (detected by PHF1 antibody). The human tissue was from 12 individuals ranging in age from 64-95 years old and with Braak stages 0-VI (Supplementary Table 1; [14]). The mouse brains were from 24-month-old CVN mice, which overexpress human APP with 3 amyloidogenic mutations, lack a nitric oxide 2 synthase gene, and accumulate plaques and tangles in an age-dependent manner [15].

The *in vivo* localization results were intended to serve as a pilot survey of the cellular and intracellular distribution of tau_pT217_, rather than a comprehensive cataloging of tau_pT217_ localization as functions of disease status and neuroanatomical location in humans or mice.

As illustrated in Supplementary Figure 5 for human brain, tau_pT217_ and tau_pS396/pS404_ were barely detectable, if at all, at Braak stage 0, although some tau_pT181_ and tau_pT231_ were evident. In contrast, all 4 phospho-tau epitopes were detectable at Braak stages I-VI, as shown in Figures 1–3, respectively. Throughout those disease stages, tau_pT217_ was primarily detected in axons, neuropil threads, and neuronal cell bodies and nuclei, and co-localized extensively with structures labeled by the pan-tau Tau5 antibody. In general, tau_pT217_ immunoreactivity was much more pronounced at Braak stages III-VI than at Braak stages I-II, and co-localization of tau_pT217_ was much more extensive with the Tau5 epitope than with tau_pT181_, tau_pT231_ or tau_pS396/pS404_. In 24-month-old CVN mice, the tau_pT217_ distribution was qualitatively similar to what we observed in Braak stages III-VI human brains in terms of labeled structures, staining intensity, and comparison with structures labeled by Tau5, and antibodies to tau_pT181_, tau_pT231_ and tau_pS396/pS404_ (Supplementary Figure 6).

### 3.3 Tau_pT217_ is associated with developing post-synaptic sites in cultured neurons

As shown in Supplementary Figures 2 and 3, under basal conditions 10-14-day old, cultured mouse cortical neurons express some tau_pT217_ that appears as puncta within MAP2-positive dendrites. Mature synapse formation is still underway in cultured neurons of that age, and to test the hypothesis that at least some of the puncta correspond to developing dendritic spines, we double-labeled such neurons with antibodies to tau_pT217_ and the dendritic spine protein, PSD95. When the cells were imaged by a spinning disc laser confocal microscopy with a resolution of ~200 nm in the x-y plane (Figure 4A, B), tau_pT217_ appeared to be extensively co-localized with PSD95, with a Pearson correlation coefficient of ~0.7 (Figure 4C).

To gain deeper insight into the spatial relation of tau_pT217_ to dendritic spines marked by anti-PSD95, we also examined doubly labeled cells using a dSTORM super-resolution microscope with an x-y resolution of ~15-20 nm (Figure 4B). Under those conditions, the Pearson correlation coefficient was only 0.1 (Figure 4C), although some pixels were positive for both proteins. When combined with the analogous data for confocal microscopy, the dSTORM data indicated that tau_pT217_ is typically located close to dendritic spines, rather than frequently being spatially coincident with them.

Further evidence for association of tau_pT217_ with dendritic spines was obtained by first defining each contiguous set of tau_pT217_-positive pixels and PSD95-positive pixels as a single object. Then, for each tau_pT217_-positive object we plotted the shortest distance to a PSD95-positive object, and conversely, the shortest distance from each PSD95-positive object to a tau_pT217_-positive object. As can be seen in Figure 4D, both types of measurements revealed that most PSD95-positive and tau_pT217_-positive objects are within 1 μm of each other, with many sharing overlapping pixels and thereby are separated by no more than 15-20 nm. Altogether, these data indicate a very close association of tau_pT217_ with developing dendritic spines.

### 3.4 Exposure of cultured neurons to xcTauOs, but not to xcAβOs, increases intraneuronal tau_pT217_

xcTauOs and xcAβOs are toxic to neurons and self-propagate in AD brain by prion-like mechanisms [19]. To seek conditions that modulate levels of intracellular tau_pT217_, we exposed cultured mouse cortical neurons to xcTauOs made from recombinant 2N4R human tau and separately to xcAβOs made from synthetic Aβ_1-42_ (Supplementary Figure 7). The total tau and Aβ_1-42_ concentrations were 250 nM and 2 μM, respectively, and the treatment times were 16-18 hours for xcTauOs and 24 hours for xcAβOs.

As shown in Figure 5, xcTauOs induced an ~56% increase in tau_pT217_ as determined by quantitative immunoblotting, and an ~13% increase as measured by quantitative immunofluorescence microscopy. Statistical signficance was achieved by both methods: p = 0.0084 by immunoblotting and p = 0.0188 by immunofluorescence. In contrast, tau_pT217_ levels were unaffected by xcAβO exposure as measured by either immunoblotting or immunofluorescence.

### 3.5 Phosphorylation at T217 reduces tau’s affinity for cytoplasmic microtubules

Tau was originally described as microtubule-associated protein in brain [20], and its sub-molecular domains that facilitate tau-tau binding in paired helical filaments [21–24] are embedded within the microtubule-binding repeat region of tau [25]. Conditions that reduce tau’s affinity for microtubules might therefore increase tau’s propensity to form toxic aggregates.

To test the hypothesis that phosphorylation at T217 reduces tau’s affinity for microtubules, we expressed EGFP-human 2N4R tau in CV-1 (African green monkey kidney) cells and performed FRAP microscopy to study the kinetics of tau turnover on microtubules. Two forms of EGFP-tau with respect to the tau sequence were expressed for FRAP: WT and pseudo-phosphorylated T217E (Supplementary Figure 8). As shown in Figure 6A, cells with microtubule-bound EGFP-tau were photographed before and immediately after photobleaching of selective regions of cytoplasm, and at several additional post-bleach times.

Loss of fluorescence in the photobleached areas simply reflected quenching of the EGFP signal, rather than a loss of tau from microtubules. This phenomenon is demonstrated by photobleaching of Venus-Tau-Teal: 2N4R human tau tagged at its N-terminus with Venus fluorescent protein and at its C-terminus with Teal fluorescent protein [17]. As can be seen in Figure 6B, following complete photobleaching of Venus, microtubules labeled by fluorescent tau can still be clearly seen in the blue-shifted Teal channel that was minimally excited by the photobleaching laser.

As illustrated in Figures 6C-E, the peaks of fluorescence recovery were achieved within ~10 seconds for the T217E forms of EGFP-tau, but the fluorescence intensity of the WT form rose continuously for 100 seconds post-bleaching, and never reached a peak. These results indicate that tau_pT217_ cycles on and off of microtubules much more rapidly than WT tau.

To verify that the T217E mutation structurally mimics tau_pT217_, we used western blotting with anti-tau_pT217_ of CV-1 cells transfected to express EGFP-tau_WT_, EGFP-tau_T217E_ or EGFP-tau_T217A_, the latter of which cannot be phosphorylated at position 217 because of the alanine mutation at that site. As shown in Supplementary Figure 8, anti-tau_pT217_ recognized EGFP-tau_WT_ and EGFP-tau_T217E_, but not EGFP-tau_T217A_. The immunoreactivity with EGFP-tau_WT_ reflects a baseline level of WT tau phosphorylation of T217. More importantly, the robust immunoreactivity of anti-tau_pT217_ with EGFP-tau_T217E_ and the absence of immunoreactivity with EGFP-tau_T217A_ demonstrates that the T217E mutation structurally mimics phosphorylation of T217. Altogether, the EGFP-tau expression experiments strongly imply that phosphorylation at T217 substantially reduces tau’s affinity for microtubules.

## 4 DISCUSSION

A flurry of recent studies of potential AD biomarkers has provided evidence that assays for CSF or blood plasma levels of tau_pT217_ can rival or exceed the accuracy of assays for other fluid biomarkers or PET imaging for detecting AD, especially when combined with other diagnostic assays [7, 9, 26]. Indeed, a study that was reported at the [Tau 2022 conference](https://alz.org/tau/overview.asp) but has not been published yet prompted a tentative conclusion that tau_pT217_ levels in CSF define a “clock” that predicts AD progression over a 30 year time span (P-Tau217 Clock). Despite these encouraging advances about the diagnostic value of tau_pT217_ for AD, to the best of our knowledge only one peer-reviewed paper [12] and one published pre-print [13] have addressed the histopathology of tau_pT217_, and not a single study published until now has investigated what tau_pT217_ does to neurons or what provokes its intracellular accumulation.

To shed more light on the pathobiology of tau_pT217_ we therefore completed a histological survey of its cellular and subcellular distribution in the brains of human AD patients and an AD model mouse strain, and of its cell biological properties in cultured neurons and fibroblasts. Doing so required optimization of an anti-tau_pT217_ antibody that can recognize other proteins with the identical epitope, an issue that likely applies to other antibodies directed against tau_pT217_. Consistent with prior immunohistological studies of human brain [12, 13], we found that tau_pT217_ first appears in limbic structure neurons at Braak stages I-II, that its presence in the somatodendritic compartment and neuropil threads increases with disease progression, and that its cellular and subcellular distribution overlaps partially with those of other phospho-tau variants associated with AD and other tauopathies. An immunofluorescence study of aged CVN mice indicates that the tau_pT217_ distribution in this particular AD model mouse strain is similar to what is seen in mid to late-stage human AD, highlighting the potential utility of CVN mice for further studies of tau_pT217_. Using cultured mouse neurons, we found baseline levels of tau_pT217_ closely associated with nascent dendritic spines, and stimulation of the intracellular tau_pT217_ level by neuronal exposure to xcTauOs, but not to xcAβOs. Finally FRAP analysis of EGFP-tagged tau in fibroblasts implied that phosphorylation of tau at T217 potently decreases tau’s affinity for microtubules. Altogether, these new data raise the possibility that phosphorylation at T217 provokes rapid tau turnover on microtubules, concomitant impairment of fast axonal transport [27] and synaptic activity, and stimulation of tau aggregation [28, 29].

Because AD biomarker assays that rely on antibodies to tau_pT217_ are being developed for clinical use, it is essential that such antibodies do not cross-react with other proteins. It is possible that the polyclonal rabbit anti-tau_pT217_ that we used for this study represents the only anti-tau_pT217_ antibody that is currently available from commercial sources, although not all vendors provide sufficient information to make be certain. The was sold by Abcam and it was produced by immunizing a rabbit with an LPpTPP peptide that is found in at least 18 human and/or mouse proteins other than tau. The possibility that this antibody, as well as the other commercially available rabbit polyclonals and proprietary antibodies to tau_pT217_, recognize one or more proteins besides tau phosphorylated at T217 is thus a serious concern. Fortunately, we found that diluting the antigen affinity purified antibody 5000-fold to a concentration of 200 ng/ml (1.43 nM) for western blotting of human brain, mouse brain and mouse neuron cultures allowed detection of tau_pT217_ without detecting other bands that were recognized by the antibody when it was used at higher concentrations (Supplementary Figures 1 and 4). We therefore also diluted the antibody to 200 ng/ml for immunofluorescence detection of tau_pT217_. This concentration-dependent specificity of the antibody presumably reflects protein-specific differences in amino acid sequences that flank the LPpTPP sequence and affect the affinity of the antibody for tau_pT217_ versus other proteins that contain the same peptide. This cautionary tale about the anti-tau_pT217_ antibody that we used should be taken into account for other basic and clinical research studies, as well as clinical applications aimed at detecting tau_pT217_.

Our immunohistological analysis of tau_pT217_ in human brain yielded results similar to those reported earlier by two other groups [12, 13]. At Braak stage 0 we did not detect any tau_pT217_ (Supplementary Figure 5), but occasional immunostaining of axons, neuropil threads and neuronal cell bodies was evident by Braak stages I-II (Figure 1), and more widespread by Braak stages III-VI (Figures 2 and 3). In light of prior reports of tau_pT217_ abundance in tangles [12, 13] we were surprised to find minimal co-localization of structures labeled by anti-tau_pT217_ and the tangle marker antibody, PHF1. One possible explanation for this discrepancy is the concentration at which anti-tau_pT217_ was used. Moloney and colleagues found tau_pT217_ at all Braak stages using a ThermoFisher rabbit polyclonal antibody that was diluted 1:500 [13] and is described by the supplier as “produced against a chemically synthesized phosphopeptide derived from the region of human tau that contains threonine 217”. Based on that description, the ThermoFisher and Abcam anti-tau_pT217_ antibodies might be identical, in which case tangle detection might depend on using the antibody for immunohistochemistry at a concentration too high to be specific for tau_pT217_. In fact, we frequently observed tangles in human AD neurons when the Abcam anti-tau_pT217_ was used at a 1:500 dilution.

Frequent tangles containing tau_pT217_ were also reported by Wennström and colleagues, who used a proprietary rabbit anti-tau_pT217_ made by Eli Lilly and Company [12]. Wennström and colleagues also reported that the Lilly antibody robustly labeled granulovacuolar degeneration bodies and multi-vesicular bodies, neither of which we observed. Because further specifications about that antibody are not publicly available, we do not know if it detected tangles and the other structures because it was used at too high a concentration, or if other factors, such as antigen retrieval methods, other aspects of the immunohistochemical protocol or neuroanatomical considerations might explain the discrepancies with our results.

To gain further insight into functions and regulation of tau_pT217_ we examined its localization and induction in cultured neurons, and used FRAP microscopy to study its association with microtubules in fibroblasts expressing EGFP-tau. Using conventional confocal and dSTORM super-resolution microscopy, we found a striking correspondence between tau_pT217_ puncta and nascent dendritic spines marked by anti-PSD95 in primary mouse cortical neurons cultured under basal conditions (Figure 4). In addition, exposure of such cultured neurons to xcTauOs caused a substantial increase in intracellular tau_pT217_, whereas xcAβOs did not affect the level of intraneuronal tau_pT217_ (Figure 5). Finally, we found that tau-EGFP pseudo-phosphorylated at position 217 by a T-to-E mutation had a dramatically shorter dwell time on fibroblast microtubules than WT tau coupled to EGFP (Figure 6). Altogether, these results raise the possibilities that tau_pT217_ is involved in synapse formation and function, that soluble xcTauOs in the brain parenchyma provoke tau_pT217_ accumulation in AD neurons *in vivo*, and that the reduced microtubule affinity of tau phosphorylated at T217 causes dysregulated fast axonal transport, which we described previously as occurring in cultured neurons derived from tau knockout mice [27]. While much further work is required to refine understanding of what causes tau_pT217_ to accumulate in neurons and how it affects neuronal functions, the evidence shown here represents a first step.

## Supporting information

all supplementary figures and tables

## ACKNOWLEDGEMENTS

This work was supported by the Owen’s Family Foundation (GSB), the Cure Alzheimer’s Fund, the Rick Sharp Alzheimer’s Foundation and NIH grant RF1 AG051085 to G.S.B. We would like to thank Dr. Dora Bigler-Wang for handling mice; Dr. Swapnil Sonkusare for help with dSTORM microscopy; Dr. Ammasi Periasamy, Director of the University of Virginia’s W.M. Keck Center for Cellular Imaging, for assistance with FRAP microscopy; NIH equipment grant OD025156 for funding purchase of the Keck Center’s Zeiss 980 confocal microscopy system; and the late Drs. Lester (Skip) Binder and Peter Davies for respectively providing us with Tau5 hybridoma cells and PHF1 antibody.

## AUTHOR CONTRIBUTIONS

B.R. co-conceived, co-designed and performed the research, and co-analyzed the data. J.W.M. provided annotated human brain samples and assisted with interpretation of immunohistochemistry images. G.S.B. co-conceived, co-designed and supervised the research. B.R., J.W.M and G.S.B. wrote/edited the manuscript. All authors approved the manuscript.

## SUPPLEMENTARY MATERIALS

**SUPPLEMENTARY FIGURE 1** Validation of anti-tau_pT217_ specificity by western blotting. Anti-tau_pT217_ and total tau (Tau5) western blots were performed against (A) cultured wild type (WT) and tau knockout (TKO) mouse cortical neurons, (B) 24 month old WT and CVN mouse brain homogenates, and (C) elderly, age-matched, cognitively normal and AD human brain homogenates. When used at 200 ng/ml (1.43 nM), as for these experiments, anti-tau_pT217_ labeled a single band with the same electrophoretic mobility as total tau, and the band was sensitive to alkaline phosphatase treatment prior to the primary antibody step and was undetectable in TKO mouse brain.

**SUPPLEMENTARY FIGURE 2** Validation of anti-tau_pT217_ specificity by immunofluorescence. Braak stage V-VI human brain (A), 24 month old CVN mouse brain (B) and cultured WT mouse cortical neurons were double-labeled with anti-tau_pT217_ at 200 ng/ml (1.43 nM), and to mark the neuronal somatodendritic compartment, with anti-MAP2. Note that anti-tau_pT217_ was sensitive to prior treatment of the tissue sections or cells with alkaline phosphatase.

**SUPPLEMENTARY FIGURE 3** tau_pT217_ is partly co-localized with tau_pT181_, tau_pT231_ and tau_pS396/pS404_ in WT mouse cortical neurons cultured under basal conditions. Neurons were obtained from E16-E17 embryonic brain and were cultured for 10-14 days. Single plane confocal images are shown.

**SUPPLEMENTARY FIGURE 4** Validation of anti-tau_pT181_ and mouse anti-tau_pT231_ antibodies in 10-14 day old primary WT and TKO mouse cortical neurons. Tau5 was used as a total tau marker.

**SUPPLEMENTARY FIGURE 5** tau_pT217_ localization in Braak stage 0 human brain cortex. Representative fields of view show ~5 μm thick sections that were triple-labeled with antibodies to tau_pT217_, MAP2, and tau_pT181_, tau_pT231_ or tau_pS396/pS404_, and imaged confocally with a 40X objective. All images illustrate single confocal planes.

**SUPPLEMENTARY FIGURE 6** tau_pT217_ is partly colocalized with tau_pT181_ and tau_pT231_ in 24 month old CVN mouse brain sections. Single plane confocal images are shown.

**SUPPLEMENTARY FIGURE 7** Analysis of extracellular oligomers of 2N4R human tau and Aβ_1-42_ by western blotting. Tau5 was used to probe the tau blot and 6E10 was used to probe the Aβ_1-42_ blot.

**SUPPLEMENTARY FIGURE 8** Characterization of EGFP-tau variants used for fluorescence recovery after photobleaching (FRAP) microscopy. The fluorescent fusion proteins were expressed in CV-1 cells as WT, T217E pseudo-phosphorylated and T217A non-phosphorylatable versions of 2N4R human tau. Note that by western blotting anti-tau_pT217_ labeled EGFP-tau_WT_ and EGFP-tau_T217E_, but not EGFP-tau_T217A_, indicating that the T217E mutation accurately mimics that structure of the tau_pT217_ epitope.

**SUPPLEMENTARY TABLE 1** Clinical characterization of human autopsy brain samples used in the study.

**SUPPLEMENTARY TABLE 2** Primary and secondary antibodies used in this study.

**SUPPLEMENTARY TABLE 3** Human and mouse proteins containing the LPpTPP sequence (from PhosphoSite Plus: https://www.phosphosite.org/homeAction).

