## Supplementary material for "Localization, induction, and cellular effects of tau-phospho-threonine 217": all supplementary figures and tables

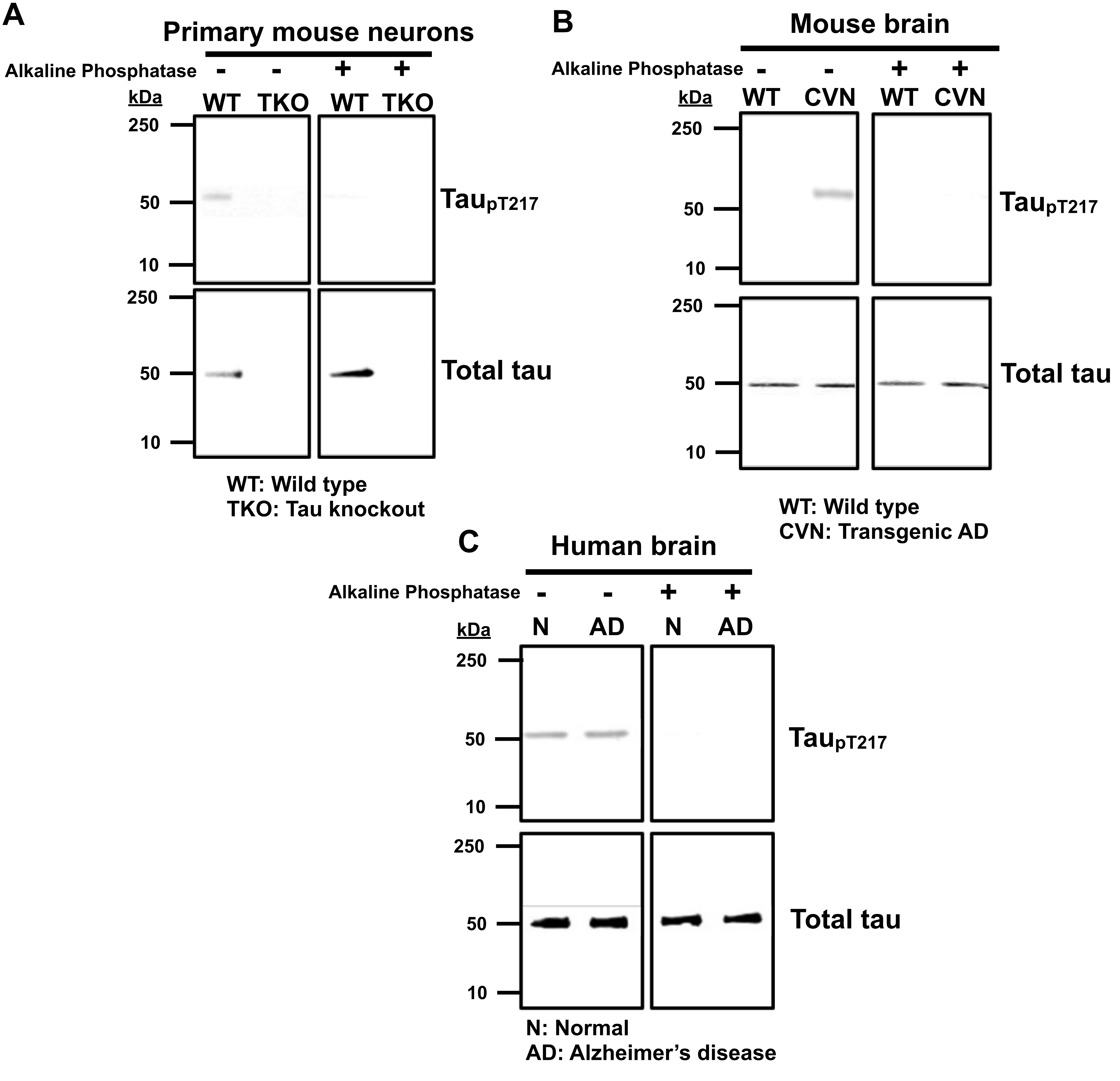


**Supplementary Figure 1** Validation of anti-tau$_{pT217}$ specificity bhy western blotting. Anti-tau$_{pT217}$ and total tau (Tau5) western blots were performed against (A) cultured wild type (WT) and tau knockout (TKO) mouse cortical neurons, (B) 24 month old WT and CVN mouse brain homogenates, and (C) elderly, age-matched, cognitively normal and AD human brain homogenates. When used at 200 ng/ml (1.43 nM), as for these experiments, anti-tau$_{pT217}$ labeled a single band with the same electrophoretic mobility as total tau, and the band was sensitive to alkaline phosphatase treatment prior to the primary antibody step and was undetectable in TKO mouse brain.

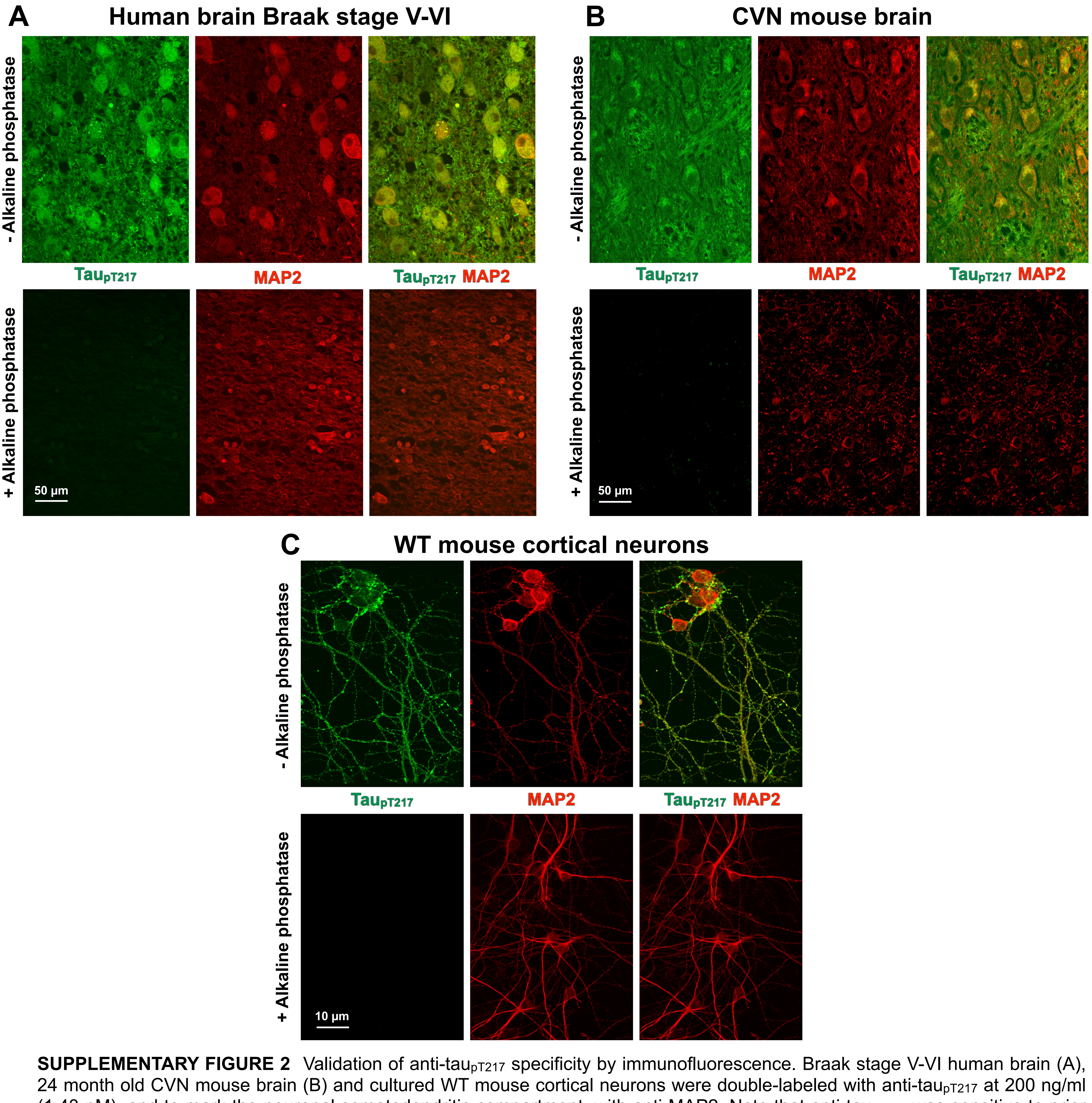


**SUPPLEMENTARY FIGURE 2** Validation of anti-tau$_{pT217}$ specificity by immunofluorescence. Braak stage V-VI human brain (A), 24 month old CVN mouse brain (B) and cultured WT mouse cortical neurons were double-labeled with anti-tau$_{pT217}$ at 200 ng/ml (1.43 nM), and to mark the neuronal somatodendritic compartment, with anti-MAP2. Note that anti-tau$_{pT217}$ was sensitive to prior treatment of the tissue sections or cells with alkaline phosphatase.

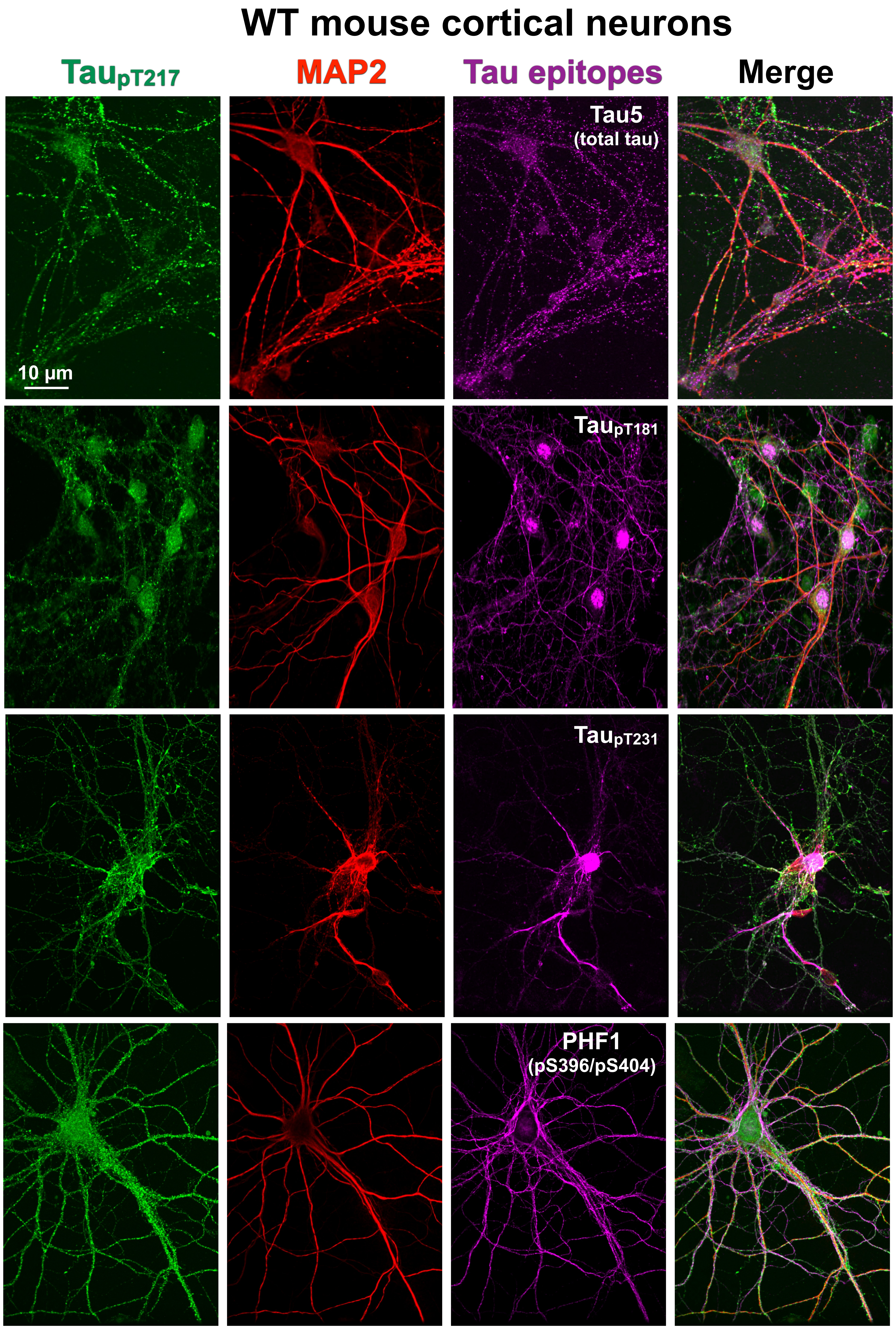


**SUPPLEMENTARY FIGURE 3** Tau$_{pT217}$ is partly co-localized with tau$_{pT181}$, tau$_{pT231}$ and tau$_{pS396/pS404}$ in WT mouse cortical neurons cultured under basal conditions. Neurons were obtained from E16-E17 embryonic brain and were cultured for 10-14 days. Single plane confocal images are shown.

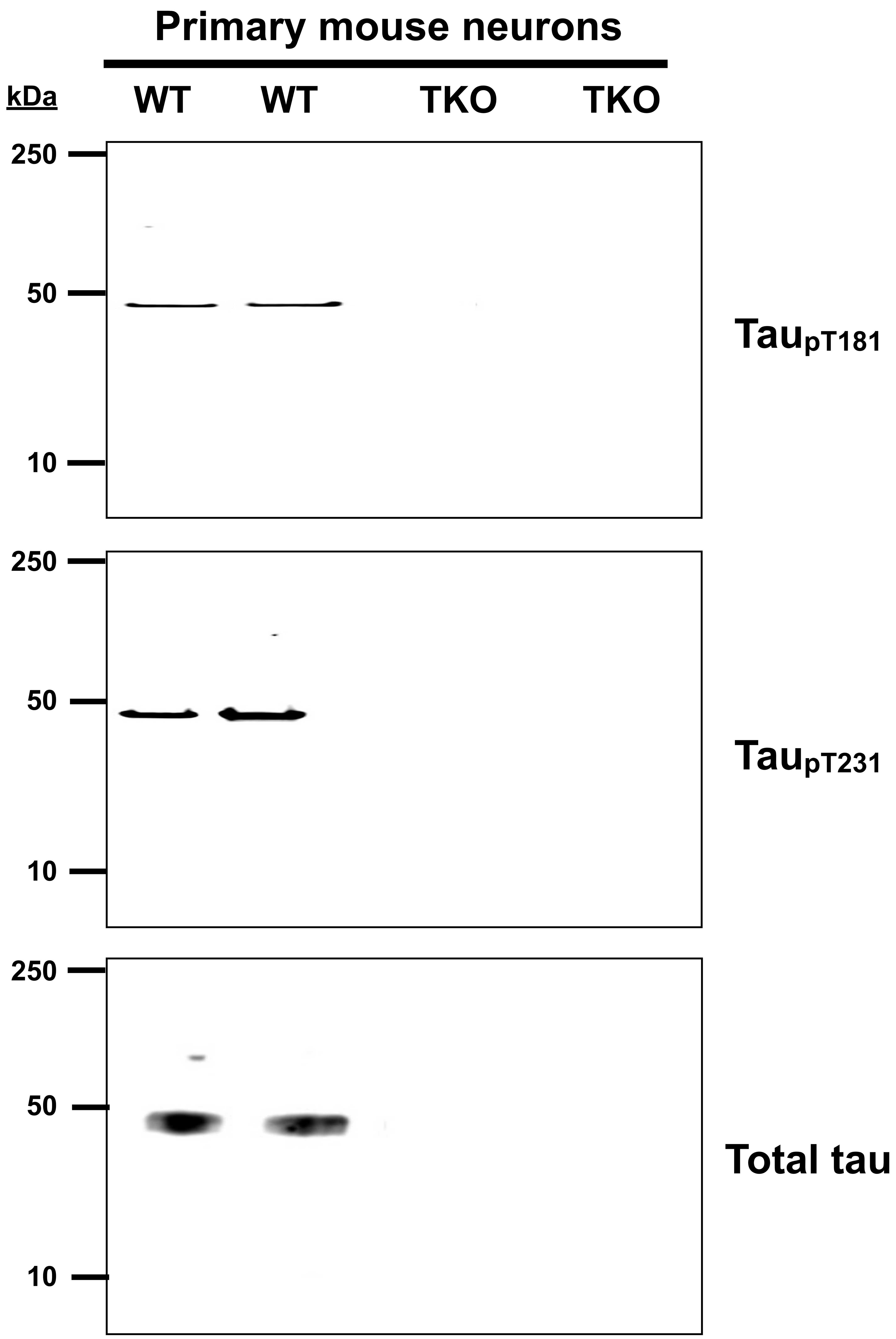


**SUPPLEMENTARY FIGURE 4** Validation of anti-tau$_{pT181}$ and mouse anti-tau$_{pT231}$ antibodies in 10-14 day old primary WT and TKO mouse cortical neurons. Tau5 was used as a total tau marker.

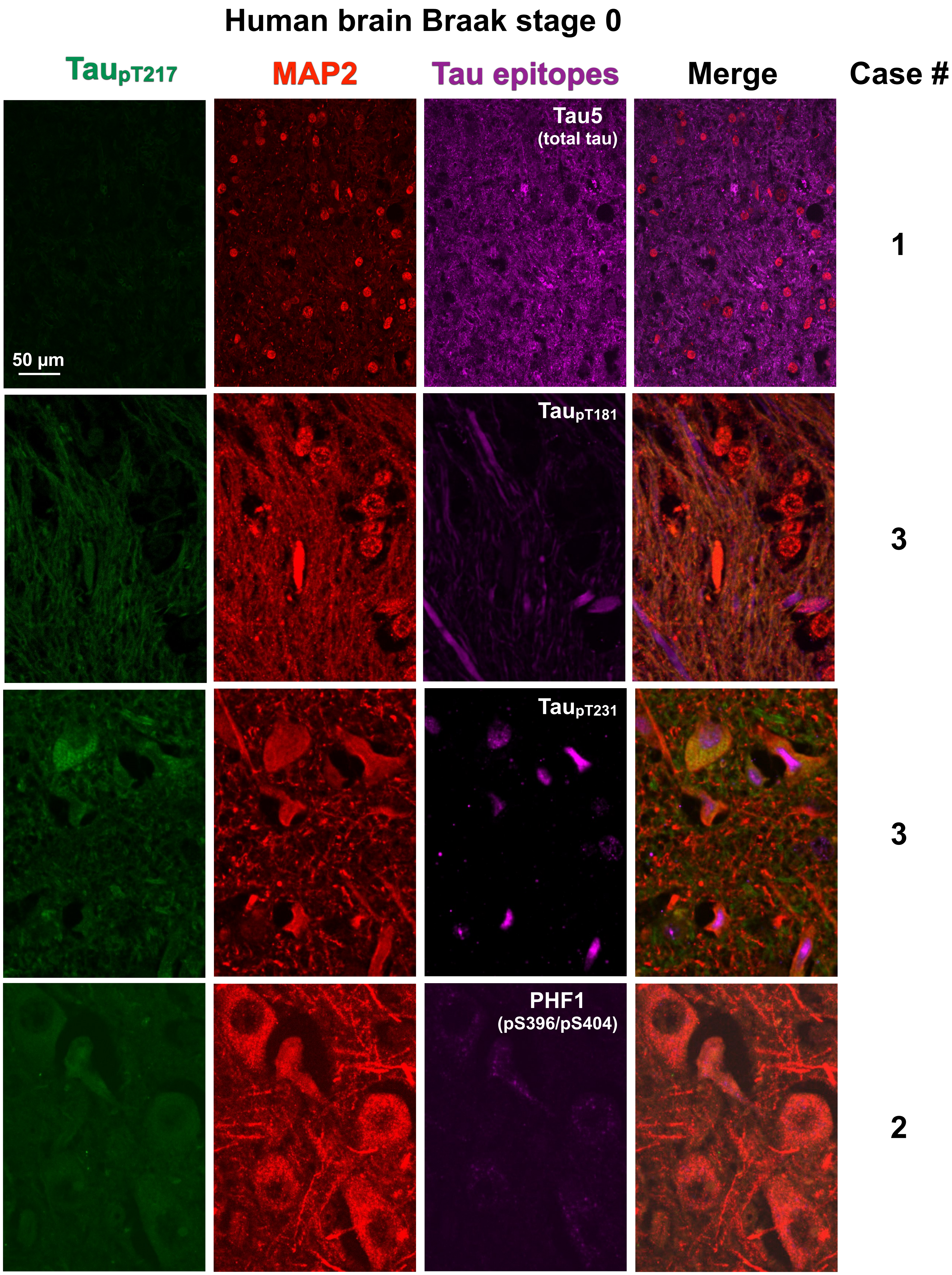


**SUPPLEMENTARY FIGURE 5** Tau$_{pT217}$ localization in Braak stage 0 human brain cortex. Representative fields of view show ~5 µm thick sections that were triple-labeled with antibodies to tau$_{pT217}$, MAP2, and tau$_{pT181}$, tau$_{pT231}$ or tau$_{pS396/pS404}$, and imaged confocally with a 40X objective. All images illustrate single confocal planes.

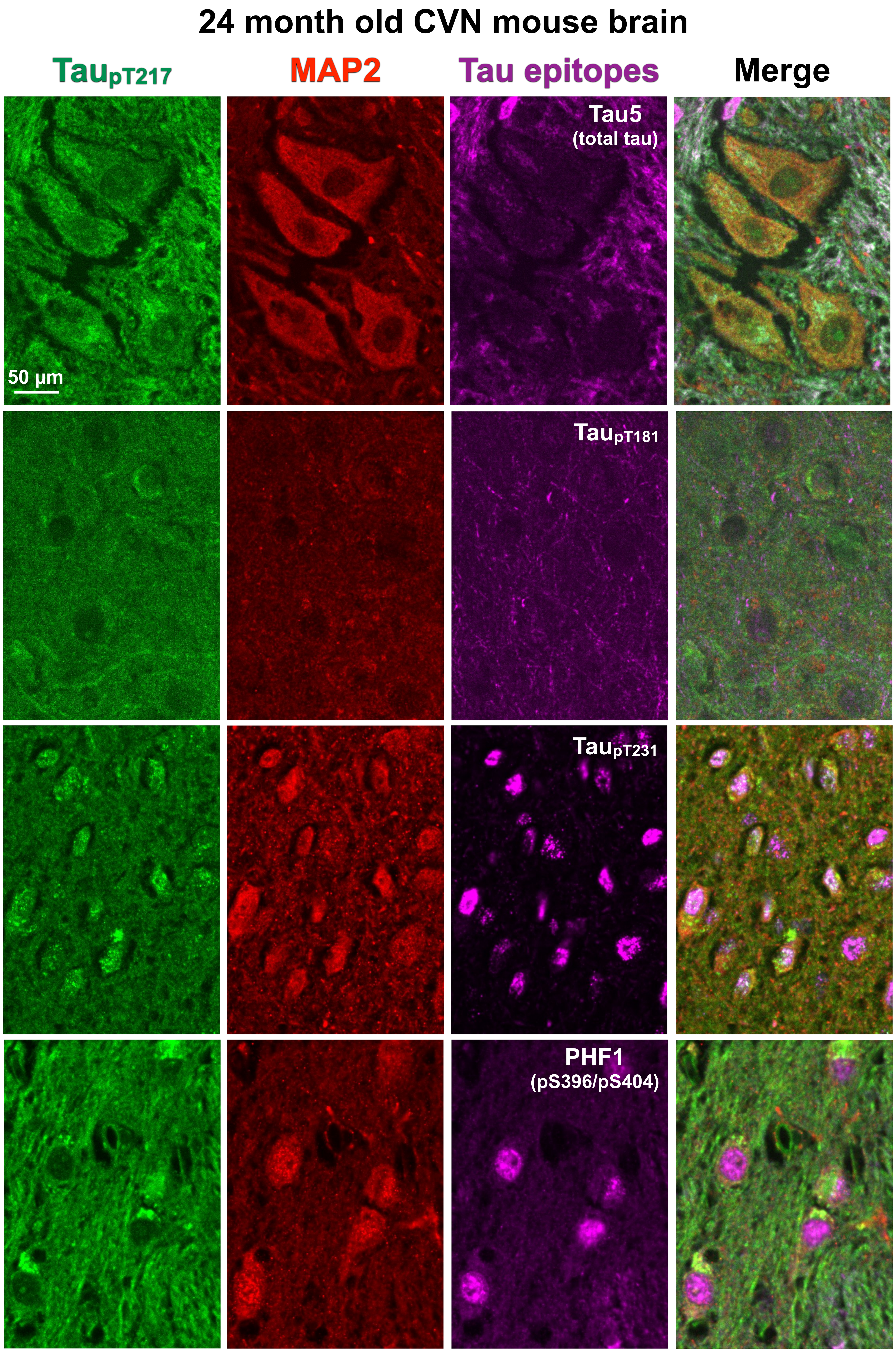


**SUPPLEMENTARY FIGURE 6** Tau$_{pT217}$ is partly colocalized with tau$_{pT181}$ and tau$_{pT231}$ in 24 month old CVN mouse brain sections. Single plane confocal images are shown.

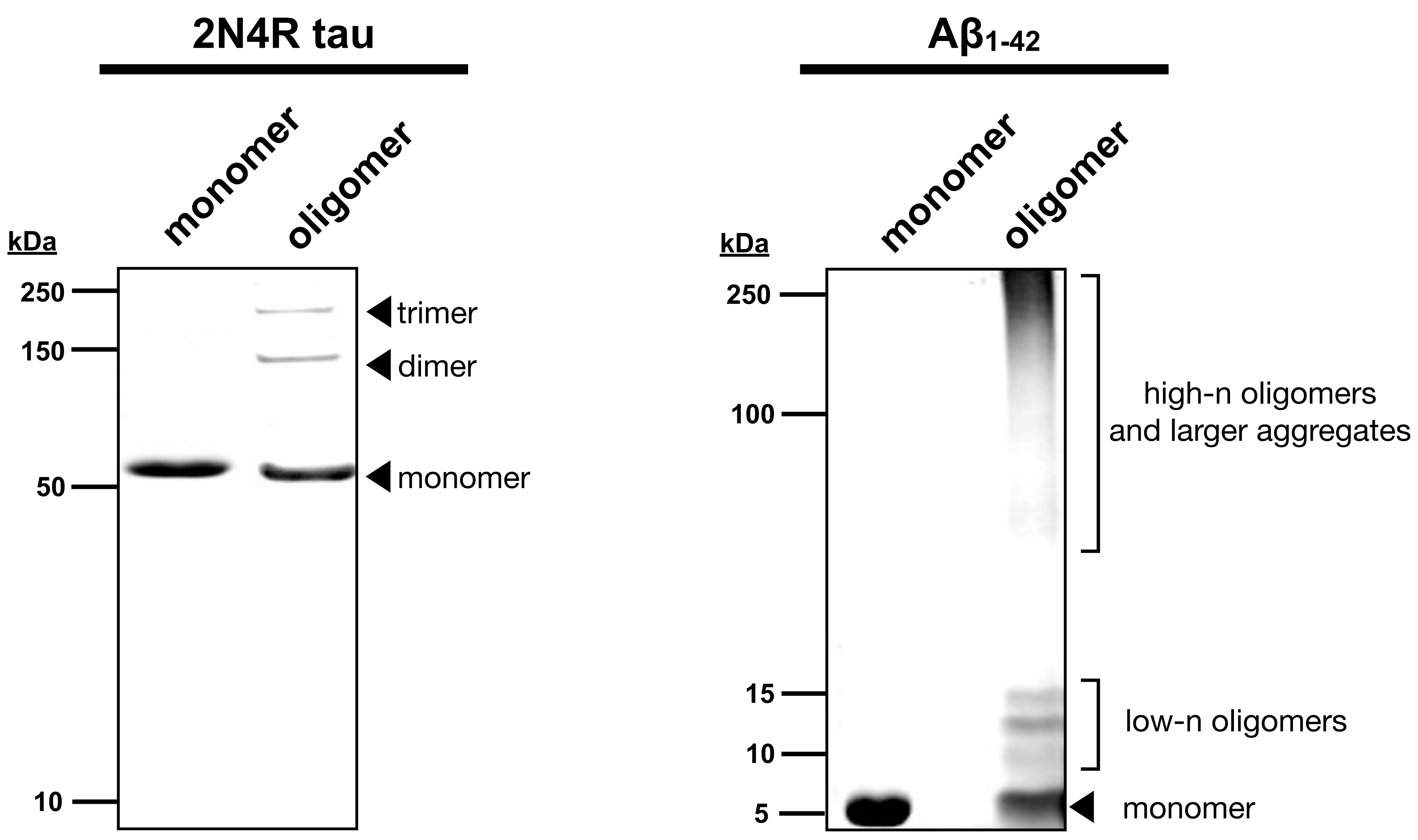


**SUPPLEMENTARY FIGURE 7** Analysis of extracellular oligomers of 2N4R human tau and A$\beta_{1-42}$ by western blotting. Tau5 was used to probe the tau blot and 6E10 was used to probe the A$\beta_{1-42}$ blot.

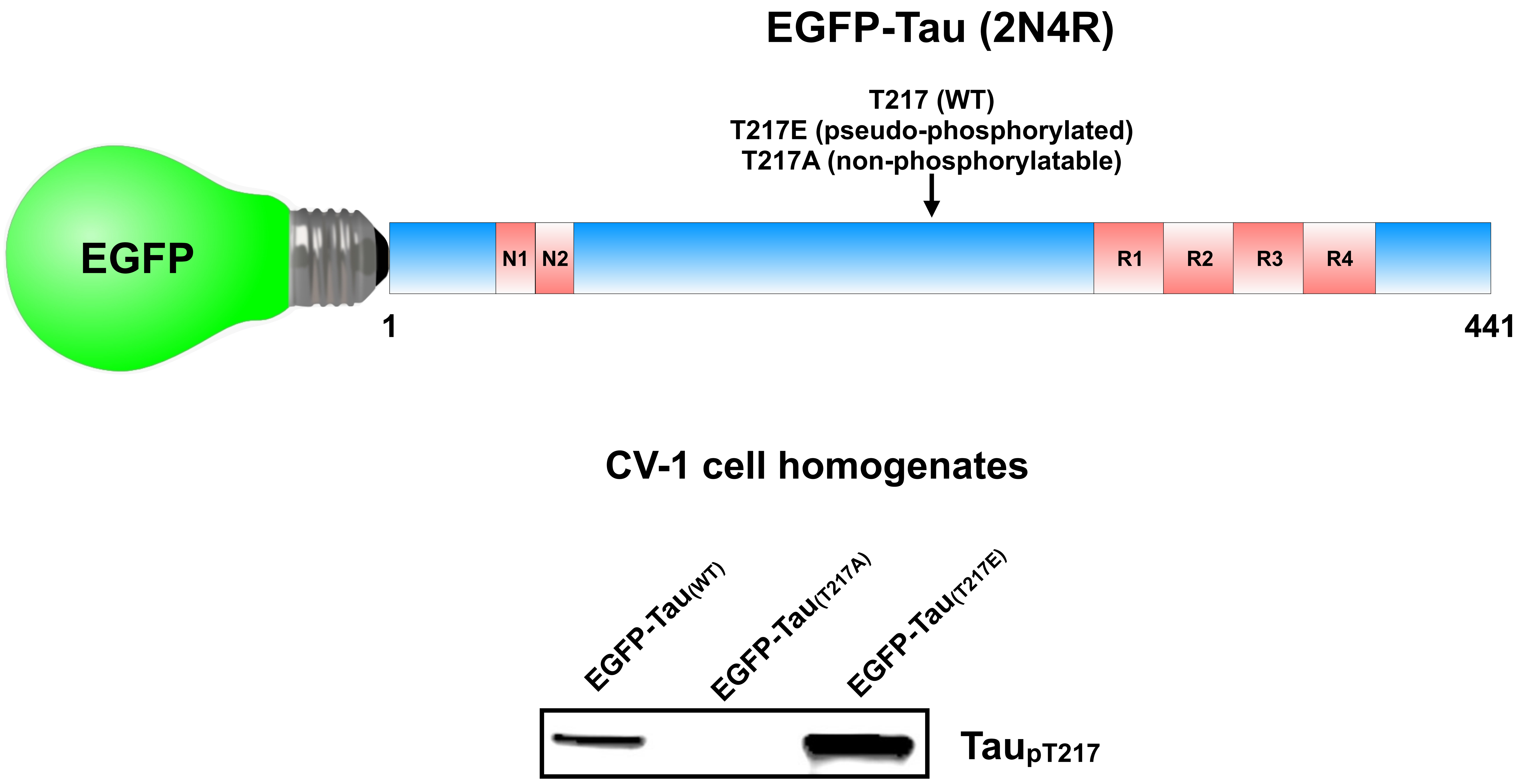


**SUPPLEMENTARY FIGURE 8** Characterization of EGFP-tau variants used for fluorescence recovery after photobleaching (FRAP) microscopy. The fluorescent fusion proteins were expressed in CV-1 cells as WT, T217E pseudo-phosphorylated and T217A non-phosphorylatable versions of 2N4R human tau. Note that by western blotting anti- tau$_{pT217}$ labeled EGFP-tau$_{WT}$ and EGFP-tau$_{T217E}$, but not EGFP-tau$_{T217A}$, indicating that the T217E mutation accurately mimics that structure of the tau$_{pT217}$ epitope.

**SUPPLEMENTARY TABLE 1. Clinical characterization of human autopsy brain samples used in the study.**

| Case # | Diagnosis | Braak stage | Age (years) | Gender | Race | Caregiver of | Postmortem delay (hours) |
|---|---|---|---|---|---|---|---|
| 1 | NML | 0 | NA | NA | NA | NA | NA |
| 2 | HTN vascular changes, acute ischemic injury | 0 | 81 | F | C | Yes | 24 |
| 3 | NML | 0 | 73 | M | C | NA | NA |
| 4 | NML | 0 | 64 | F | C | NA | NA |
| 5 | FTD/MND TDP subtype and early AD A2B1C2 | I-II | 68 | M | C | NA | NA |
| 6 | early AD | I-II | 82 | M | C | NA | NA |
| 7 | AD A3B3C3 and CAA | V-VI | 93 | F | C | NA | 24 |
| 8 | AD A2B2C2 | III-IV | 88 | F | C | NA | 48 |
| 9 | AD A3B2C3 | III-IV | 79 | M | C | NA | NA |
| 10 | AD A1B3C3 | V-VI | 89 | M | C | NA | NA |
| 11 | AD A2B2C2 CAA | III-IV | 95 | M | C | NA | 24 |
| 12 | AD A3B3C3 and CAA | V-VI | 95 | F | C | NA | NA |

**Abbreviations:**

NML = Normal

AD = Alzheimer’s disease

HTN = Hypertension

FTD/MND = Frontotemporal dementia with motor neuron disease

CAA = Cerebral amyloid angiopathy

**SUPPLEMENTARY TABLE 2. Primary and secondary antibodies**

| Name | Tag | Source/Cat no. |
|---|---|---|
| Mouse anti-pan-tau (Tau5) | None | Lester (Skip) Binder (deceased) |
| Rabbit anti-tau$_{pT217}$ | None | Abcam/ab192665 |
| Mouse anti-tau$_{pT231}$ | None | BioLegend/828901 |
| Mouse anti-tau$_{pT181}$ | None | BioLegend/846602 |
| Chicken anti-MAP2 | None | Abcam/ab92434 |
| Mouse anti-GFAP | None | Thermo Fisher Scientific/MA5-12023 |
| Mouse anti-BIN1 (clone 99D) | None | Millipore/05-449 |
| Mouse anti-PSD95 | None | MyBioSource/MBS804156 |
| Mouse anti-PHF-tau (PHF1) | None | Peter Davies (deceased) |
| Mouse anti-amyloid-β (6E10) | None | BioLegend/803001 |
| Goat anti-rabbit IgG | Alexa Fluor 488 | Invitrogen/A11034 |
| Goat anti-mouse IgG | Alexa Fluor 488 | Invitrogen/A11029 |
| Goat anti-chicken Igγ | Alexa Fluor 568 | Invitrogen/A11041 |
| Goat anti-mouse IgG | Alexa Fluor 647 | Invitrogen/A21235 |
| Goat anti-rabbit IgG | Alexa Fluor 647 | Invitrogen/A21244 |
| Goat anti-rabbit IgG | IRDye 680 | LI-COR/926-68071 |
| Goat anti-mouse IgG | IRDye 800 | LI-COR/926-32210 |

**SUPPLEMENTARY TABLE 3. Human and mouse proteins containing the LPpTPP sequence**
**(from PhosphoSite Plus: https://www.phosphosite.org/homeAction)**

| GENE | PROTEIN | SPECIES | ACCESSION # | PROTEIN MOLECULAR WEIGHT | AMINO ACID START POSITION | AMINO ACID END POSITION | MODIFICATION SITE (relative to *human or #mouse 2N4R tau) |
|---|---|---|---|---|---|---|---|
| *ATOH8* | ATOH8 | mouse | UP:Q99NA2 | 34,785 | 123 | 127 | T125 |
| *CEPT1* | CEPT1 | human<br>mouse | UP:Q9Y6K0<br>UP:Q8BGS7 | 46,554<br>46,434 | 38 | 42 | T40 |
| *DIDO1* | DATF1 | human | UP:Q9BTC0 | 243,873 | 1660 | 1664 | T1662 |
| *KMT2C* | MLL3 | human<br>mouse | UP:Q8NEZ4<br>UP:Q8BRH4 | 541,370<br>540,187 | 3975<br>3968 | 3979<br>3972 | T3977<br>T3970 |
| *KMT2D* | MLL4 | human<br>mouse | UP:O14686<br>UP:Q6PDK2 | 593,389<br>600,245 | 4680<br>4731 | 4684<br>4735 | T4682<br>T4733 |
| *MYC* | Myc | human<br>mouse | UP:P01106<br>UP:P01108 | 48,804<br>48,971 | 56 | 60 | T58 |
| *MYCN* | N-Myc | human<br>mouse | UP:P04198<br>UP:P03966 | 49,561<br>49,572 | 56 | 60 | T58 |
| *MAP3K14* | Nik | mouse | UP:Q9WUL6 | 103,080 | 688 | 692 | T690 |
| *ZNF746* | PARIS | human | UP:Q6NUN9 | 69,136 | 601 | 605 | T603 |
| *PEX12* | PEX12 | human | UP:O00623 | 40,797 | 279 | 283 | T281 |
| *PHACTR3* | PHACTR3 | human<br>mouse | UP:Q96KR7<br>UP:Q8BYK5 | 62,552<br>62,652 | 234<br>233 | 238<br>237 | T236<br>T235 |
| *POU4F1* | POU4F1 | human | UP:Q01851 | 42,697 | 37 | 41 | T39 |
| *PPIP5K2* | PPIP5K2 | human | UP:O43314 | 140,407 | 1190 | 1194 | T1192 |
| *Rab11fip5* | RAB11FIP5 isoform 2 | mouse | UP:Q6ZQ33 | 142,909 | 1126 | 1130 | T1128 |
| *RALGPS1* | RALGPS1 | human<br>mouse | UP:Q5JS13<br>UP:A2AR50 | 62,133<br>65475 | 329 | 333 | T331 |
| *RAVER1* | RAVER1 isoform 2 | human<br>mouse | UP:Q8IY67-2<br>UP:Q9CW46 | 77,860<br>79,382 | 486<br>492 | 490<br>496 | T488<br>T494 |
| *ESRP1* | RBM35A | human<br>mouse | UP:Q6NXG1<br>UP:Q6NXG1 | 75,585<br>75,549 | 424<br>423 | 428<br>427 | T426<br>T425 |
| *MAPT* | Tau | human<br>mouse | UP:P10636<br>UP:P10637 | 36,760 - 78,928<br>35,714 - 76,243 | *215<br>#204 | *219<br>#208 | *T217<br>#T206 |
| *VARS1* | VARS | human | UP:P26640 | 140,476 | 282 | 286 | T284 |
